# Assessment of the Potential Role of *Streptomyces* in Cave Moonmilk Formation

**DOI:** 10.1101/131847

**Authors:** Marta Maciejewska, Delphine Adam, Aymeric Naômé, Loïc Martinet, Elodie Tenconi, Magdalena Całusińska, Philippe Delfosse, Marc Hanikenne, Denis Baurain, Philippe Compère, Monique Carnol, Hazel Barton, Sébastien Rigali

**Affiliations:** InBioS - Centre for Protein Engineering, Institut de Chimie B6a, University of Liège, B-4000, Liège, Belgium; Environmental Research and Innovation Department, Luxembourg Institute of Science and Technology, Rue du Brill 41, Belvaux, L-4422, Luxembourg; InBioS - Functional Genomics and Plant Molecular Imaging, University of Liège, B-4000 Liège, Belgium; PhytoSYSTEMS, University of Liège, B-4000 Liège, Belgium; InBioS – Eukaryotic Phylogenomics, University of Liège, B-4000, Liège, Belgium; Department of Biology, Ecology and Evolution & Centre of Aid for Research and Education in Microscopy (CAREm-ULg), Institute of Chemistry B6a University of Liège, B-4000, Liège, Belgium; InBioS - Plant and Microbial Ecology, Botany B22, University of Liège, B-4000, Liège, Belgium; Department of Biology, University of Akron, Akron, Ohio, United States of America

**Keywords:** biomineralization, moonmilk genesis, geomicrobiology, carbonatogenesis, cave microbiology

## Abstract

Moonmilk is a karstic speleothem mainly composed of fine calcium carbonate crystals (CaCO_3_) with different textures ranging from pasty to hard, in which the contribution of biotic rock-building processes is presumed to involve indigenous microorganisms. The real bacterial input in the genesis of moonmilk is difficult to assess leading to controversial hypotheses explaining the origins and the mechanisms (biotic versus abiotic) involved. In this work we undertook a comprehensive approach in order to assess the potential role of filamentous bacteria, particularly a collection of moonmilk-originating *Streptomyces*, in the genesis of this speleothem. Scanning electron microscopy (SEM) confirmed that indigenous filamentous bacteria could indeed participate in moonmilk development by serving as nucleation sites for CaCO_3_ deposition. The metabolic activities involved in CaCO_3_ transformation were furthermore assessed *in vitro* among the collection of moonmilk *Streptomyces*, which revealed that peptides/amino acids ammonification, and to a lesser extend ureolysis, could be privileged metabolic pathways participating in carbonate precipitation by increasing the pH of the bacterial environment. Additionally, *in silico* search for the genes involved in biomineralization processes including ureolysis, dissimilatory nitrate reduction to ammonia, active calcium ion transport, and reversible hydration of CO_2_ allowed to identify genetic predispositions for carbonate precipitation in *Streptomyces*. Finally, their biomineralization abilities were confirmed by environmental SEM, which allowed to visualize the formation of abundant mineral deposits under laboratory conditions. Overall, our study provides novel evidences that filamentous Actinobacteria could be key protagonists in the genesis of moonmilk through a wide spectrum of biomineralization processes.

## Introduction

The hypogean environment, although highly deprived of nutrients, sustains a diverse microbial life. This subterranean microbiome plays an important ecological role in caves, with secondary effects on mineralogy, including host rock dissolution or mineral precipitation, leading to the formation of various secondary mineral deposits termed speleothems (Barton and Northup 2007; Jones 2010; Cuezva et al. 2012). A biogenic origin has been hypothesized for a number of speleothems, including coralloids (Banks et al. 2010), pool fingers (Melim et al. 2001), ferromanganese deposits (Northup et al. 2003; Spilde et al. 2005), helictites (Tisato et al. 2015), and moonmilk (Cañaveras et al. 2006). Unlike typical mineral deposits, moonmilk is present as a soft and pasty precipitate on cave surfaces and within pools (Richter et al. 2008; Cacchio et al. 2014). The origin of moonmilk has been controversial for many years due to the complex mineralogy, the atypical crystalline morphology, and also the size of its crystals (Verrecchia and Verrecchia 1994; Cañaveras et al. 1999; Cañaveras et al. 2006; Bindschedler et al. 2010; Bindschedler et al. 2014). Notably, moonmilk deposits are characterized by several crystal habits including nano-fibers, and micro-meter sized needle-fiber crystals in a form of monocrystalline rods and polycrystalline chains (Cañaveras et al. 1999; Cañaveras et al. 2006; Bindschedler et al. 2010; Bindschedler et al. 2014). While initially postulated as a speleothem of abiotic origin (Harmon et al. 1983; Borsato et al. 2000), recent studies attributed the genesis of moonmilk to indigenous microbial population (Cañaveras et al. 2006; Cailleau et al. 2009; Baskar et al. 2011; Braissant et al. 2012).

Microbial carbonate precipitation (MCP) is a broad spectrum phenomenon either mediated by autotrophic pathways, such as photosynthesis and methanogenesis that lead to depletion of local CO_2_, or heterotrophic pathways that alter local conditions to promote CaCO_3_ precipitation (Castanier et al. 2000; Banks et al. 2010). Caves are devoid of sunlight, ruling out photosynthesis, while methanogenesis has been documented rarely in these systems. Heterotrophic processes may therefore play an important role (Banks et al. 2010). Active calcite precipitation by heterotrophs in calcium-rich environments has been hypothesized to be the consequence of a detoxification process, wherein the acidification of the local environment induced by passive influx of Ca requires growing cells that would actively export the excess of this metal to maintain cellular calcium homeostasis (Banks et al. 2010). Such metal detoxification strategies have also been linked to the formation of the unusual speleothems known as helictites (Tisato et al. 2015). Heterotrophic growth can also increase the environmental pH, which can in turn increase the saturation index of CaCO_3_ and drive precipitation. Calcite precipitation through nitrogen metabolism is thought to operates by different metabolic pathways, including ureolysis, ammonification through amino acid and peptide catabolism, and dissimilatory nitrate reduction to ammonia (DNRA), all of which increase the local pH (Fig. 1) (Castanier et al. 2000).

**Fig. 1.**
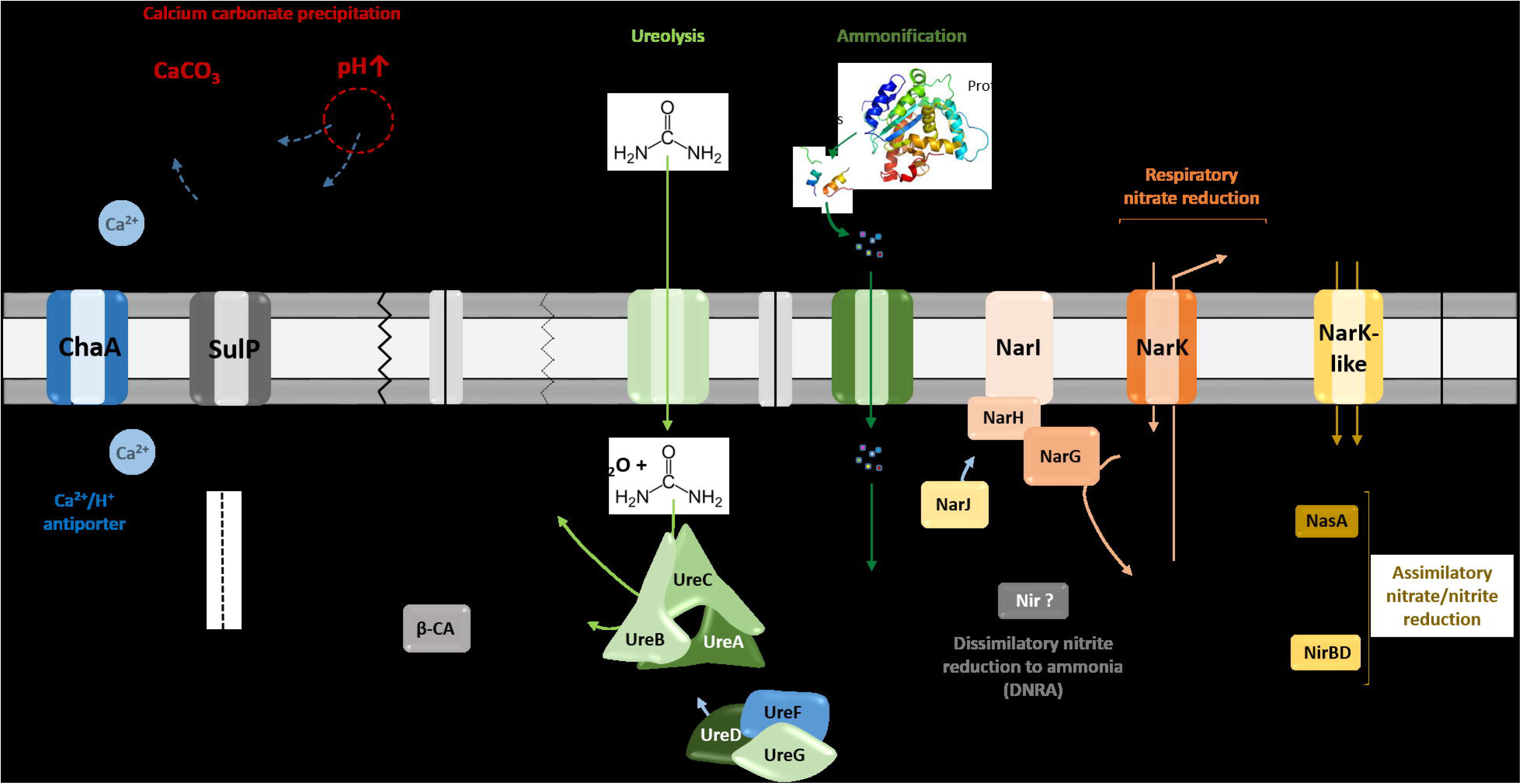
Heterotrophic pathways associated with carbonatogenesis presumed to occur in *Streptomyces* species. Microbially mediated carbonate precipitation might be either linked to active transport of calcium ions across cellular membrane through Ca^2+^/2H^+^ antiporter system – ChaA, or to nitrogen cycle-related pathways – ureolysis, ammonification and dissimilatory reduction of nitrate to ammonium/ammonia (DNRA). All those pathways lead to alkalinization of the bacterial environment through the generation of ammonium, shifting the equilibrium towards carbonate/bicarbonate ions, which, upon the presence of calcium, precipitate as calcium carbonate (CaCO_3_). ChaA, apart from providing calcium ions for potential precipitation, locally increase pH through simultaneous incorporation of protons. The urease activity seems to be linked with cytoplasmic carbonic anhydrases (β-CA), which catalyze dehydration of carbonic acid produced during ureolysis into carbon dioxide that can constitute an additional source of bicarbonate ions for precipitation. The export or import of bicarbonate ions could be potentially mediated via the sulfate transporter family protein in cluster with CA (SulP-type permease). Assimilation of organic nitrogen from amino acids releases extracellular ammonia through deamination, once the metabolic demand for nitrogen is fulfilled. The inorganic nitrogen source (NO_3_^−^), which is imported in *Streptomyces* by Nark-type transporters, can participate into biomineralization exclusively through dissimilatory reduction to ammonia mediated by respiratory nitrate reductases (Nar) together with potential nitrite reductases (Nir), which would be functional under dissimilatory conditions. Reduction of nitrite to ammonia in streptomycetes was found to operate through assimilatory pathway catalyzed by NasA and NirBD reductases

Moonmilk was reported to host wide spectrum of microbiota, including Archaea, Bacteria, and Fungi (Rooney et al. 2010; Portillo and Gonzalez 2011; Engel et al. 2013; Reitschuler et al. 2014; Reitschuler et al. 2015; Axenov-Gibanov et al. 2016; Maciejewska et al. 2016; Reitschuler et al. 2016). Among this microbiome, Fungi and filamentous microorganisms, particularly members of Actinobacteria phylum were reported to be possibly involved in the moonmilk genesis (Cañaveras et al. 2006; Bindschedler et al. 2010; Bindschedler et al. 2014). Bindschedler et al. (2010, 2014) suggested that the presence of nano-fibers within the crystalline structure of moonmilk was associated with biomineralized fungal hyphae. The authors suggest that organized networks of nano-fibers, often observed in moonmilk, could represent fibrous fungal cell wall polymers, such as chitin and β-(1→3)glucans (Bindschedler et al. 2010). On the other hand, the observation of unstructured aggregates of unconsolidated microcrystalline fibers with calcified Actinobacteria-like filaments led Cañaveras et al. (2006) to propose a model of moonmilk formation wherein Actinobacteria promoted calcium carbonate precipitation by creating locally favorable conditions, with the bacterial cell walls serving as nucleation zones (Cañaveras et al. 2006). The presence of metabolically active microorganisms in moonmilk was demonstrated using isothermal microcalorimetry (Braissant et al. 2012), although the progressive accumulation of CaCO_3_ (and presumably entombment) ultimately leads to a decrease of the microbial activity (Cañaveras et al. 2006; Sanchez-Moral et al. 2012). As a result, microorganisms would play a significant role in the initiation of moonmilk formation, which ultimately would be overtaken by abiotic processes leading to the growth of the deposit, which can reach up to 1 m in thickness (Sanchez-Moral et al. 2012).

In all of these studies, there has been no clear distinction as to whether the increase or decrease in local pH is ultimately leading to the precipitation of moonmilk by the dominant actinobacterial species observed. In this work we use a combination of microscopy, cultivation and genomic approaches to provide an *in vitro* and *in silico* assessment of the actinobacterial metabolic activities that could promote CaCO_3_ precipitation. Our data suggest that the *Streptomyces* species would play an important role in nitrogen metabolism, which could locally raise pH and contribute to moonmilk formation.

## Materials and methods

### Moonmilk sampling and *Streptomyces* strains used in this study

Samples for microscopy and cultivation were taken from moonmilk deposits originating from three sampling points (collection points COL1, COL3, COL4, supplementary Fig. 1 from Maciejewska et al. 2016) in the upper Viséan limestone cave ‘Grotte des Collemboles’ (Springtails’ Cave), Comblain-au-Pont, Belgium (more detailed cave description in supplementary Fig. 1). Moonmilk samples for strains isolation were brought to the laboratory on ice and stored at 4°C prior to lyophilization. Extensive attempts at the cultivation of Actinobacteria led to the isolation of 31 phylogenetically distinct *Streptomyces* strains representing phylogenetically distinct phylotypes (as previously described in Maciejewska et al. 2016 and supplementary Fig. 2). Moonmilk samples for scanning electron microscopy (SEM) were preserved in two separate fixative solutions, 2.5% glutaraldehyde in 0.1 M Na-acetate buffer (pH 7.4), and 100% ethanol, and stored at 4°C until analysis. To exclude the effect of fixatives on the crystalline structure of moonmilk, the lyophilized samples were also observed under the microscope as controls (data not shown).

### Environmental scanning electron microscopy (ESEM) and elemental X-ray energy dispersive microanalysis (EDS)

Glutaraldehyde-fixed moonmilk samples were post-fixed in 1% osmium tetroxide (OsO_4_) in distilled water, rinsed and dehydrated through a graded ethanol series (30-100%). Glutaraldehyde-fixed samples and ethanol-preserved samples were then processed by critical point drying, prior to mounting on the glass slides covered by double-side carbon tape together with lyophilized samples. Parts of each sample were mounted to expose the outer surface, a vertical section or fracture made with a scalpel blade. Dry samples were subsequently sputter coated with platinum on Balzers sputtering Unit SCD 030 (Balzers, Lichtenstein).

The production of mineral deposits by isolates MM24 and MM99 following growth in two different culture conditions - calcite precipitation agar (CPA) and modified B-4 medium (more details in CaCO_3_ precipitation section) was detected from the living bacterial colonies after being air-dried. Morphological observations were performed by light microscopy (reflected, transmitted and polarized light) using an Olympus Provis AX-70 microscope fitted with a Visicam 5.0 videocamera for image capture. SEM observations were performed in an environmental scanning electron microscope FEI XL30 ESEM-FEG (Eindhoven, The Netherlands). Platinum-coated samples were observed under high vacuum (HV) conditions with the ET-secondary electron (SE) detector at 10 mm working distance and 15 kV accelerating voltage. Air-dried cultures, were observed under low vacuum (LV) conditions (0.4 Torr) with the large field gaseous secondary electron (GSE) detector and the backscattered electron (BSE) detector at 10 mm working distance and 10 and 20 kV accelerating voltage, respectively. As a result of applying two types of detection – GSE and BSE, the contrast of the images due to the surface morphology or due to the atomic number of the elements and the density of minerals could be obtained. Elemental X-ray microanalysis and mapping was carried out using a Bruker silicon drift energy dispersive detector (SDD Quantax 129 eV, Billerica, MA, USA) at 10-20 kV accelerating voltage with the Esprit 1.9 software. Semi-quantitative analyses of the elemental composition were done using the standard-less ZAF method with automatic background subtraction.

### Genomic analysis of moonmilk-derived isolates

The genes of interest (see Fig. 1) including those coding for i) the ureolytic system (*ure*), ii) the Ca^2+^/2H^+^ antiporter system (*chaA*), iii) the nitrate/nitrite reductases (*nar/nas/nir*) with the corresponding transporters (*narK*), and iv) the carbonic anhydrase (CA) together with sulfate transporter in cluster with CA (*sulP*) (Felce and Saier 2005), were retrieved from the genomes of moonmilk *Streptomyces* sequenced at the Luxembourg Institute of Science and Technology, as previously described (Maciejewska et al. 2016). These genes were first identified within the chromosome of the model *Streptomyces* species *– Streptomyces coelicolor* (Bentley et al. 2002). The designations of the selected genes are listed in supplementary Table 1. Subsequently, genes sequences encoding the corresponding proteins were identified within the genomes of additional 54 reference *Streptomyces* strains (supplementary Table 2) for which completely assembled genomes are available in NCBI FTP server (data retrieved on January 8^th^ 2016).

A total of 407,461 protein sequences were organized in clusters of orthologous groups (COGs) using Proteinortho v 5.12 (Lechner et al. 2010) with the PoFF extension to further discriminate similar sequences based on synteny. Created COGs were used as models to screen moonmilk *Streptomyces* genomes. For every *S. coelicolor* gene, the collection of protein sequences in the corresponding COG was used to build a hidden Markov model (HMM) profile (Eddy 1998). As an example, the gamma sub-unit of the urease metallo-protein of *S. coelicolor* (UreA, SCO1236) clusters in a COG with 52 sequences from other *Streptomyces* (supplementary Table 1). This COG is used to construct a HMM profile representing the UreA protein that is used to search a database of translated predicted coding sequences of the moonmilk *Streptomyces*. Partial coding sequences resulting from the fragmented nature of the moonmilk *Streptomyces* genomes were also considered in the screening. The moonmilk *Streptomyces* coding sequences were predicted with Prodigal v2.6.2 (Hyatt et al. 2010). HMM profile building and HMM search were carried out using the HMMER3 software package (v3.1b2, http://hmmer.org/). The accession numbers of genes recovered from moonmilk *Streptomyces* are compiled in the supplementary Table 3.

## Metabolic assays

### Ammonification

The ability of isolates to decompose organic nitrogen into ammonia was tested on nutrient agar containing: peptone, 5 g/l; beef extract, 3 g/l; NaCl, 5 g/l; phenol red, 0.012 g/l; agar, 15 g/l; pH 7.0 (Food and Agriculture Organization of the United Nations 1983). Each representative moonmilk *Streptomyces* isolate was spot-inoculated on an individual Petri dish and incubated for 7 days at 28°C. *Citrobacter freundii* ATCC 43864 was used as a positive control strain, while uninoculated media was used as a negative control. The development of a pink color, indicating a pH increase due to the formation of ammonia following peptides/amino acids degradation was monitored every day during the incubation.

### Ureolysis

Rapid screening of urease activity was performed on Christensen’s Urea Agar Base (UAB) solid media as described previously (Hammad et al. 2013). The UAB medium was prepared as follows: urea, 20.0 g/l; NaCl, 5.0 g/l; peptone, 1.0 g/l; glucose, 1.0 g/l; KH_2_PO_4_, 2.0 g/l; phenol red, 0.012 g/l and agar, 15.0 g/l; pH 6.5. All components of the media were autoclaved except urea which was filter-sterilized and added after autoclaving. The UAB medium without urea was used as a negative control. Both types of media were inoculated with representative of each phylotype together with urease negative control strain (*Escherichia coli* ATCC 25922) and urease positive control strain (*Klebsiella pneumoniae* ATCC 13883) and incubated at 28°C. Plates were examined continually to record development of the pink color indicating a pH increase as a result of urease enzyme activity, leading to generation of ammonia through urea degradation, according to the following reaction: (NH_2_)_2_CO + H_2_O → 2NH_3_ + CO_2._

### Nitrate and nitrite reduction

The assessment of nitrate and nitrite reduction was performed as described by Li et al. (2016), with small modifications. Briefly, the phylotypes were inoculated in nitrate or nitrite agar slants (potassium nitrate / sodium nitrite, 1g/l; peptone, 5 g/l; beef extract, 3 g/l; agar, 12 g/l; pH 7.0) and nitrate / nitrite broth tubes (potassium nitrate / sodium nitrite, 1g/l; peptone, 5 g/l; beef extract, 3 g/l; pH 7.0) equipped with the Durham tubes. *Escherichia coli* ATCC 25922 was used as a control strain for exclusive reduction of nitrate to nitrite, while *Pseudomonas aeruginosa* ATCC 27853 as a complete reducer of nitrate into the nitrogen gas, collected in the Durham tube. Uninoculated media were used as an additional control. Inoculated test tubes were incubated at 28°C for 9 days in static conditions, to reduce amount of dissolved oxygen. After incubation time few drops of sulfanilic acid and alpha-naphthylamine were added to each test tube, which together react with nitrite generating red/pink color. Reduction of nitrate (NO_3_^−^) is indicated by appearance of red color in nitrate broth, while red color disappearance in nitrite broth indicates nitrite (NO_2_^−^) reduction.

### Oxidative glucose breakdown

The glucose oxidative test was carried out according to Hugh and Leifson (1953). Briefly, each representative of the phylotypes was spot-inoculated on Hugh and Leifson’s OF basal medium, prepared as follows: peptone, 2.0 g/l; NaCl, 5.0 g/l; bromothymol blue, 0.03 g/l; K_2_HPO_4_, 0.3 g/l; agar, 3.0 g/l; pH 7.1. Filter-sterilized glucose was added after autoclaving to a final concentration of 1%. The OF medium not supplemented with glucose was used as a negative control, and oxidative *Pseudomonas aeruginosa* ATCC 27853 strain as a positive control. The inoculated plates were incubated at 28°C during 7 days and monitored continually to observe development of yellow color indicating on the acid production due to glucose metabolism.

### CaCO_3_ precipitation

The screening for isolates able to precipitate calcium carbonate through ureolysis was performed using calcite precipitation agar (CPA) as previously described (Stocks-Fischer et al. 1999). The CPA medium was prepared as follows: (g/l); nutrient broth, 3.0 g/l; urea, 20.0 g/l; CaCl_2_*2H_2_O, 28.5 g/l; NaHCO_3_, 2.12 g/l; NH_4_Cl, 10.0 g/l; Agar, 15.0 g/l. All components were autoclaved apart from urea, which was added filter-sterilized. The same medium, but without urea, was used as an additional control. Alternatively, precipitation of CaCO_3_ was tested on modified B-4 medium (pH 7.0) composed of: yeast extract, 4 g/l; calcium acetate, 2.5 g/l; agar, 15 g/l. The plates inoculated with MM24 and MM99, following incubation at 28°C, were examined under ESEM after 2 months of incubation for CPA media, and 1 month of incubation for B-4.

### CaCO_3_ solubilization

Isolates were tested for their ability to solubilize calcium carbonate on two different media including i) minimal medium (MM) (Kieser et al. 2000) containing CaCO_3_ (2 g/l), supplemented or not with glucose (5 g/l), and ii) 1:100 diluted nutrient agar (Portillo et al. 2009), containing CaCO_3_ (2.5 g/l), supplemented or not with glucose (2 g/l). The spot-inoculated plates were incubated for 4 weeks at 28°C and the capability of calcium carbonate solubilization was confirmed by observation of the clear halo around a colony.

## Results

### Microscopic evaluation of indigenous moonmilk filamentous bacteria as nucleation sites for carbonate precipitation

Filamentous bacteria, particularly Actinobacteria, have been proposed to participate in the genesis of moonmilk deposits by serving as nucleation sites for carbonate deposition (Cañaveras et al. 2006). Regardless the fixation procedure (glutaraldehyde, ethanol, and freeze-drying), classical SEM observations revealed the same crystal morphologies that have also been described in the literature (data not shown) (Cañaveras et al. 1999; Cañaveras et al. 2006; Bindschedler et al. 2010; Bindschedler et al. 2014). The surface of moonmilk samples revealed the presence of dense, unstructured meshes of micrometer-size filaments known as needle-fiber calcite (Fig. 2a), which, based on EDS analysis, were shown to be mainly composed of calcium, carbon and oxygen (supplementary Fig. 3a). The structure of moonmilk deposits was characterized by the presence of abundant, randomly oriented, monocrystalline rods and polycrystalline fibers composed of stacked rhombohedra (Fig. 2a). Those crystals showed variable dimensions, ranging from 0.5 – 1 µm width and 30 - 100 µm length for monocrystals, and 2 - 20 µm width, 10 - 100 µm length for polycrystals, as previously reported (Cañaveras et al. 1999). Interestingly, we observed within the moonmilk microscopic composition, abundant networks of filaments with nano-sized width (50 - 150 nm), which either formed compacted pellets (Fig. 2b) or mats (Fig. 2c). The organized networks of nano-fibers, similar to the one observed in Fig. 2c, were previously reported from moonmilk and associated with fungal wall polymers (Bindschedler et al. 2010; Bindschedler et al. 2014). However, randomly-oriented, compacted pellets of nano-fibers as presented on Fig. 2b, were highly comparable to Actinobacteria, but characterized by a much smaller cell size which could be a result of the oligotrophic nature of the cave environment. It has been demonstrated that the cell size is largely dependent on the nutritional status of the environment, with resource-poor ecosystems stimulating dwarfism (Young 2006; Portillo et al. 2013). In this work, the observed size of the putative bacterial nano-filaments, ranging from 0.05 to 0.15 µm, could also be a consequence of the oligotrophic nature of the moonmilk niche as cell elongation/filamentation has been shown to be the result of nutritional stress in some bacteria (Steinberger et al. 2002), including Actinobacteria (Pine and Boone 1967; Wills and Chan 1978; Deutch and Perera 1992). However, even in nutrient-rich soils the majority of bacteria can display a diameter less than 0.2 µm (Hahn 2004). In addition, the imprints of nano-sized microorganisms were already reported from other geological formations, such as sedimentary rocks (Folk and Chafetz 2000; Folk 1993). Finally, recent findings of ultra-small marine Actinobacteria with an average diameter of about 0.3 µm, provides an additional evidence for the existence of nano-bacteria in oligotrophic ecosystems (Ghai et al. 2013). Unlike crystal fibers, the tiny filaments observed in glutaraldehyde- and ethanol-fixed samples, were not completely straight and regular in their shape but rather displayed plasticity and were often curved, either without preferential orientation (Fig. 2b) or in the same direction (Fig. 2d/e). The tiny, curved filaments were not observed - probably not preserved - in freeze-dried samples that suffered of ice crystal growth (data not shown). One-directional growth is not a typical behavior of growing actinobacterial filaments that are commonly branching in diverse directions in order to form a complex mycelial network. This rigid and unidirectional growth might suggest that they represent calcified filaments potentially still actively growing at their tip, which remains curved (Fig. 2d). Elemental analysis performed with EDS revealed that the observed nano-size filaments contained higher content of carbon and oxygen in comparison to the surrounding crystals, suggesting a possible biological origin (supplementary Fig. 3b). However, these EDS analyses should be viewed with great prudence as the difference in the elemental compositions between filaments and crystals could simply be a consequence of the structure and not the nature (organic vs mineral) of the analyzed areas. Additionally, along (Fig. 2f) or on the tip (Fig. 2g) of some filaments, a possible initiation of calcium carbonate deposition was observed. Besides nano-sized filaments, reticulated filaments were sporadically observed within moonmilk crystals (Fig. 2h). Those particular filamentous forms, with the size of about 0.5 µm width and up to 75 µm length, are often found in subsurface environment, including limestone or lava caves, and possess higher carbon content than the one typically observed for calcite minerals, suggesting their biogenic origin. However, the associated microorganisms are not yet identified (Melim et al. 2008; Northup et al. 2011; Miller et al. 2012; Melim et al. 2015). Overall, these observations tend to confirm the hypothesis of filamentous microorganisms (bacteria and fungi) serving as a nucleation sites for moonmilk mineral deposition (Cañaveras et al. 2006; Bindschedler et al. 2010; Bindschedler et al. 2014).

**Fig. 2.**
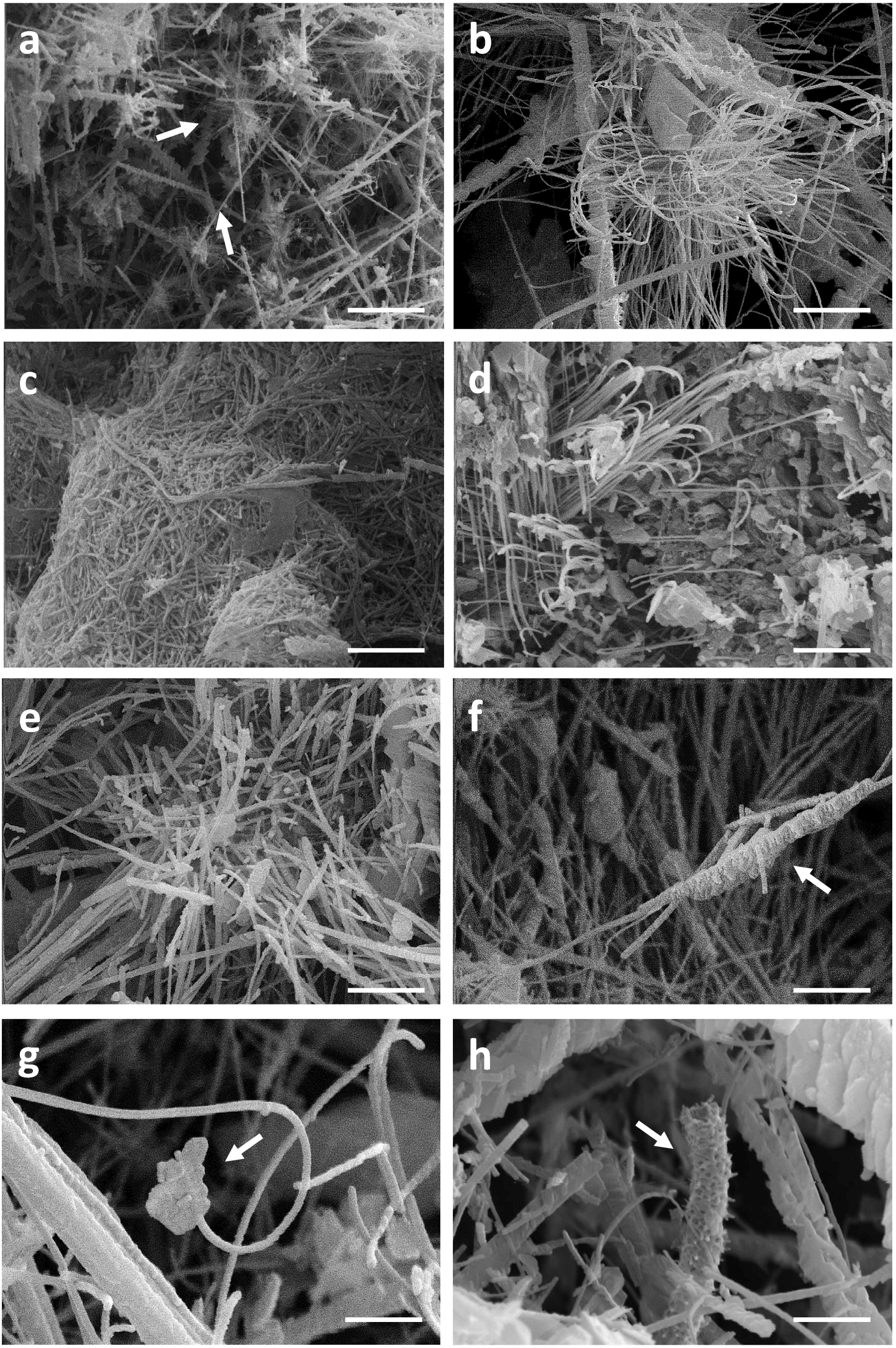
SEM-images of nano-size filaments found in the crystal structure of moonmilk deposits from the cave “Grotte des Collemboles” (Comblain-au-Pont, Belgium). Among typical moonmilk monocrystalline rods and polycrystalline fibers indicated by white arrows **(a)**, dense meshes of tiny filaments were observed, which were compacted into the stacked pellets **(b)** or dense biofilms **(c)**, mostly randomly orientated, but occasionally one-way directed **(d, e)**. Along **(f)** or on the tip **(g)** of some of those filaments, calcium carbonate deposition was identified, with some of the filaments presenting reticulated morphology **(h)** as indicated by arrows

### Potential role of cultivable moonmilk-derived *Streptomyces* in carbonate precipitation

A nucleation site itself is not sufficient to promote CaCO_3_ precipitation, as dead bacterial cells lose the ability to precipitate minerals (Banks et al. 2010). The dominant role of bacteria in calcification is attributed to metabolic activities which increase the pH of the environment (above pH 8) and therefore favor a shift of the CO_2_ – HCO_3_^−^ – CO_3_^2-^ equilibrium towards carbonate ions which precipitate with Ca^2+^ ions. We screened 31 representative *Streptomyces* strains isolated from moonmilk (see supplementary Fig. 2 retrieved from Maciejewska et al. 2016) for their metabolic activities that could lead to a raise in pH, including ureolysis, peptides/amino acids ammonification, and dissimilatory nitrate/nitrite reduction to ammonia (Fig. 1). Although these assays were performed under laboratory conditions, a qualitative assessment of these processes allowed ranking the tested phylotype representatives according to their metabolic performance, and therefore their potential to drive biomineralization through an increase in the extracellular pH. The results are shown in Fig. 3, along with a compilation of activities and phylogenetic relationships (Fig. 4). The results of each metabolic activity assay for all the tested strains are presented in supplementary Fig. 4.

**Fig. 3.**
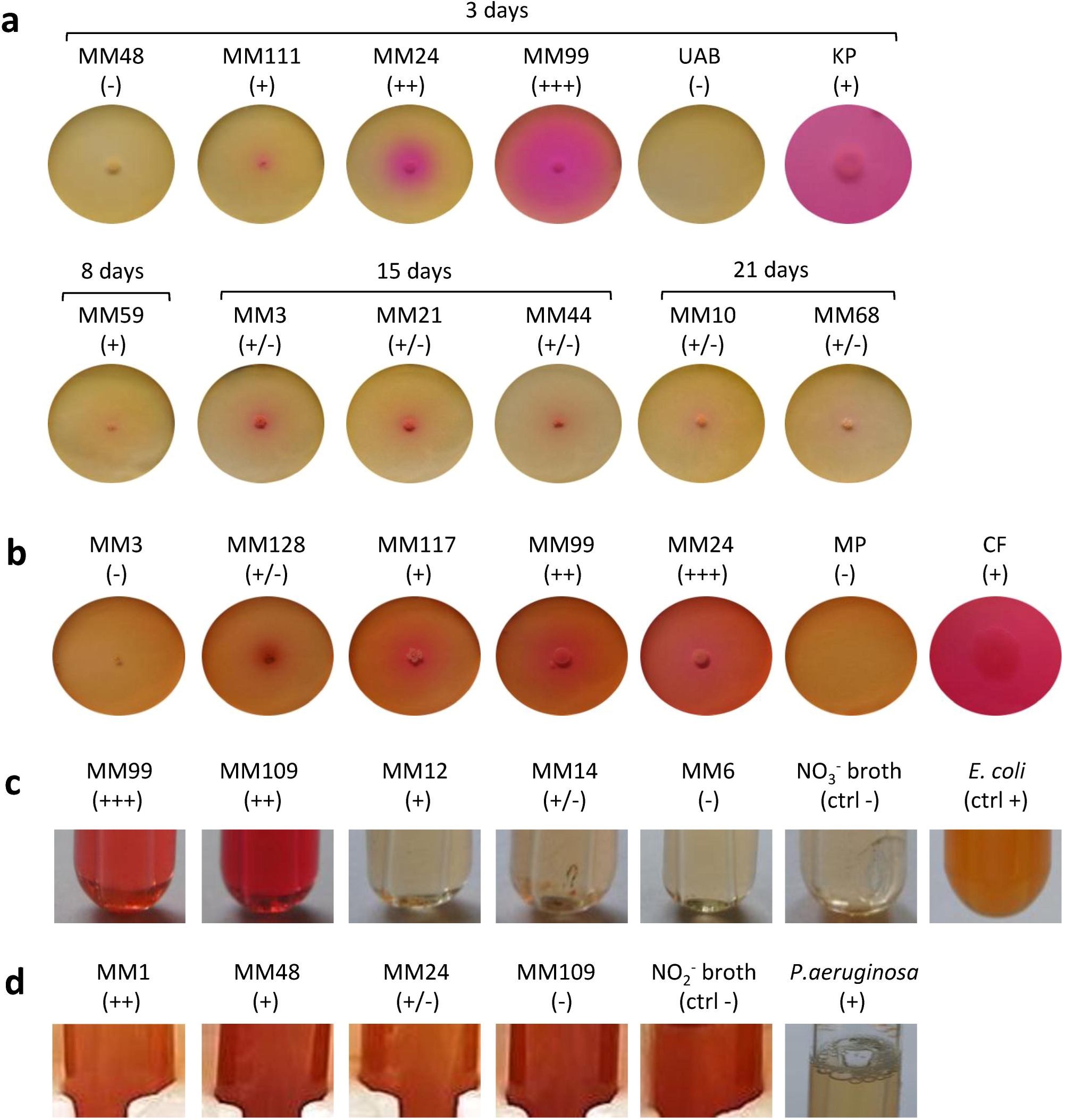
Precipitation-related metabolic activities of the moonmilk *Streptomyces*. **(a)** Ureolysis. **(b)** Ammonification. **(c)** Nitrate reduction. **(d)** Nitrite reduction. *Klebsiella pneumoniae* ATCC 13883 (KP), *Citrobacter freundii* ATCC 43864 (CF), *Escherichia coli* ATCC 25922 (EC) and *Pseudomonas aeruginosa* ATCC 27853 (PA) were used as positive controls for ureolysis, ammonification, nitrate and nitrite reduction tests, respectively. The observed activities are visualized for representative strains demonstrating different metabolic performance for each activity tested, designated through the symbols: (-) lack, (+/-) weak, (+) moderate, (++) good, and (+++) strong

**Fig. 4.**
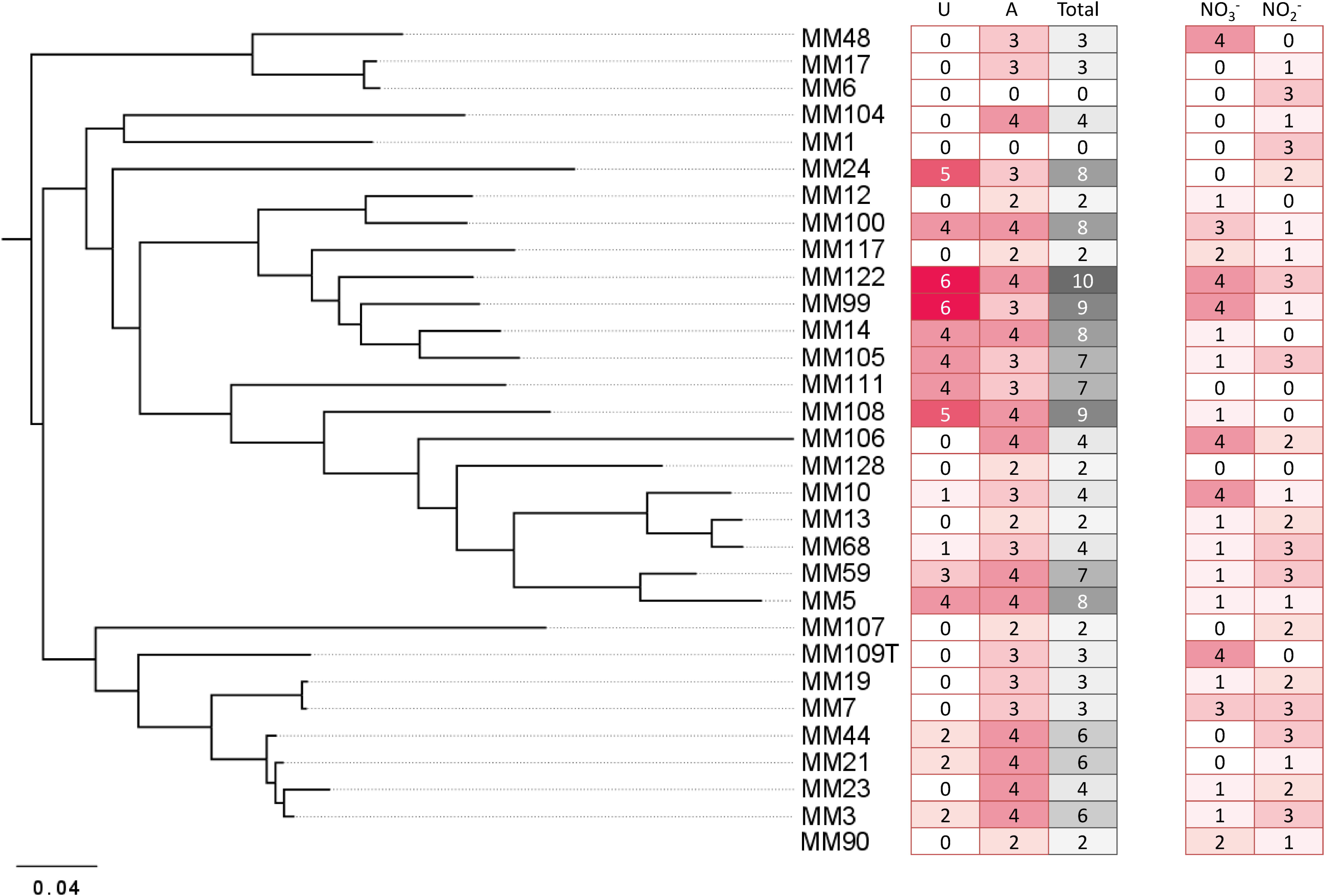
The global pattern of carbonate precipitation-associated metabolic activities of moonmilk *Streptomyces.* The heatmap plot representing metabolic performance of moonmilk *Streptomyces* strains indicated by color scale, correlated with their MLSA-based phylogenetic classification (Maciejewska et al. 2016). The total value represents the sum of metabolic performance observed for ureolysis and amino acids ammonification, ranking the isolates according to their metabolic predispositions for calcium carbonate precipitation for the tested activities. The metabolic performance of nitrate/nitrite reduction test was not included into the final ranking. Abbreviations: U, ureolysis; A, peptide/amino acid ammonification; T, total sum of activities for ureolysis and amino acid ammonification; NO_3_^−^, nitrate reduction; NO_2_^−^, nitrite reduction

### Ureolysis

Among the 31 isolates tested, 15 showed an ability to increase the pH by hydrolysis of urea (Fig. 4). MM99 and MM122 were the strongest ureolytic strains, with comparable metabolic performance to the positive control strain - *Klebsiella pneumoniae* ATCC 13883 (Fig. 3a and supplementary Fig. 4a). The majority of urease-positive moonmilk isolates displayed weak and moderate activities, which were observed either within 3 days of incubation (5 strains) or after extended time (more than 1 week) (6 strains) (Fig. 3a and supplementary Fig. 4a).

In order to know if a relation could be established between the assessed *in vitro* activity and the genetic predispositions for ureolysis, we examined the genomes of the phylotype representatives for the presence of urease genes. We used HMM profiles constructed from Clusters of Orthologous Groups (COGs) of proteins of the urease structural subunits (UreA/UreB/Ure(AB)/UreC), as well as the accessory proteins (UreF/UreG/UreD). The corresponding genes originate from functional clusters of three types *i*.*e*., *ureABCFGD, ure(AB)C*, and *ure(AB)CFGD*, which are all present in the urease-positive species. The large majority of moonmilk strains (90%) was found to encode all the urease genes, in some cases present in several copies (Table 1), suggesting that even the strains displaying urease-negative phenotypes are capable of urease activity. This suggests that in urease-negative strains urease activity is not expressed under the conditions tested, or that they harbor mutations that prevent expression or activity. Nonetheless, urea transport appears to be functional in all urease-negative isolates as urea exerted a toxic effect in the growth media of these strains (an effect that could be reversed by growth in the same medium lacking urea; data not shown).

**Table 1.** *In silico* prediction of individual genes putatively involved in the moonmilk biomineralization process retrieved from moonmilk-originating *Streptomyces* strains, including genes related to ureolysis (*ure*), nitrate/nitrite transport (*narK*) and reduction (*nar*/*nas*/*nir*), carbon dioxide (CO_2_) hydration (carbonic anhydrase - CA), and active transport of calcium ions through Ca^2+^/2H^+^ antiporter system (*chaA*).

| strain | Ureolysis |  |  |  |  |  |  | NO<sub>3</sub><sup>-</sup> / NO<sub>2</sub><sup>-</sup> transport |  | Respiratory NO<sub>3</sub><sup>-</sup> reduction |  |  |  |  | Assimilatory NO<sub>3</sub><sup>-</sup> reduction | NO<sub>2</sub><sup>-</sup> reduction |  |  | CO<sub>2</sub> hydration |  |  | Ca<sup>2+</sup> transport |
| --- | --- | --- | --- | --- | --- | --- | --- | --- | --- | --- | --- | --- | --- | --- | --- | --- | --- | --- | --- | --- | --- | --- |
|  | *ureA* | *ureB* | *ureAB* | *ureC* | *ureF* | *ureG* | *ureD* | *narK2* | *narK* | *narG* | *narH* | *narJ* | *narJ2* | *narI* | *nasA* | *nirB1* | *nirB2* | *nirD* | β-CA (1) | β-CA (2) | *sulP* | *chaA* |
| MM1 | 1 | 1 | 2 | 3 | 1 | 1 | 2 | - | + | - | - | - | - | - | 1 | 1 | 1 | 1 | 2 | 1 | 1 | - |
| MM3 | 1 | 1 | 2 | 3 | 2 | 2 | 3 | - | + | - | - | - | - | - | 1 | 1 | 1 | 1 | 2 | 1 | 1 | - |
| MM5 | 1 | 1 | 2 | 3 | 2 | 2 | 2 | - | + | - | - | - | - | - | 1 | 1 | 1 | 1 | 3 | 3 | 1 | 1 |
| MM6 | 1 | 1 | 1 | 3 | 1 | 1 | 1 | - | + | - | - | - | - | - | 1 | 1 | 2 | 2 | 4 | - | 1 | 1 |
| MM7 | 2 | 2 | 2 | 3 | 2 | 2 | 2 | + | + | 1 | 1 | - | 1 | 1 | 1 | 1 | 1 | 1 | 1 | 1 | 1 | - |
| MM10 | 1 | 1 | 1 | 2 | 2 | 2 | 2 | + | + | 3 | 2 | 1 | 1 | 2 | 1 | 1 | 1 | 1 | 2 | 1 | 1 | - |
| MM12 | 1 | 1 | 2 | 3 | 3 | 3 | 3 | - | + | - | - | - | - | - | 1 | 1 | 1 | 1 | 2 | 2 | - | 1 |
| MM13 | 1 | 1 | 1 | 2 | 2 | 2 | 2 | + | + | 1 | - | - | - | 1 | 1 | 1 | 1 | 1 | 2 | 1 | 1 | 1 |
| MM14 | 1 | 1 | 2 | 3 | 2 | 2 | 2 | - | + | - | 1 | - | 1 | 1 | 1 | 1 | 1 | 1 | 1 | 2 | 1 | - |
| MM17 | 1 | 1 | 3 | 3 | 2 | 2 | 2 | - | + | 1 | - | - | - | - | 1 | 1 | 1 | 1 | 6 | 1 | 1 | 1 |
| MM19 | 1 | 1 | 1 | 2 | 1 | 1 | 1 | - | + | - | - | - | - | - | 1 | 1 | 1 | 1 | 1 | 1 | 1 | - |
| MM21 | 1 | 1 | 2 | 3 | 2 | 2 | 2 | - | + | - | - | - | - | - | 1 | 1 | 1 | 1 | 2 | 1 | 1 | - |
| MM23 | 1 | 1 | 2 | 3 | 2 | 2 | 2 | - | + | - | - | - | - | - | 1 | 1 | 1 | 1 | 1 | 1 | 1 | - |
| MM24 | 2 | 2 | 2 | 5 | 2 | 2 | 4 | + | + | - | - | - | - | - | 1 | 1 | 1 | 1 | 2 | 2 | 1 | 1 |
| MM44 | 1 | 1 | 2 | 3 | 2 | 2 | 2 | - | + | - | - | - | - | - | 1 | 1 | 1 | 1 | 2 | 1 | 1 | - |
| MM48 | 2 | 2 | 2 | 4 | 2 | 2 | 2 | - | + | 1 | 1 | - | 1 | 1 | 2 | 1 | 2 | 1 | 5 | 1 | 1 | 1 |
| MM59 | 1 | 1 | - | 1 | 1 | 1 | 1 | - | + | - | - | - | - | - | 1 | 1 | 1 | 1 | 3 | 2 | 1 | 1 |
| MM68 | 1 | 1 | 1 | 2 | 3 | 2 | 2 | - | + | - | - | - | - | - | 1 | 1 | 1 | 1 | 2 | 2 | 1 | 1* |
| MM99 | 3 | 3 | 1 | 4 | 3 | 3 | 3 | + | - | - | - | - | - | - | 1 | - | - | - | 1 | 1 | 1 | - |
| MM100 | 1 | 1 | 2 | 4 | 2 | 2 | 2 | + | + | - | - | - | - | - | 2 | 1 | 1 | 1 | 2 | 1 | - | - |
| MM104 | 1 | 1 | 1 | 1 | 1 | 1 | 1 | - | + | - | - | - | - | - | - | 1 | 1 | 1 | 2 | 3 | 1 | 1 |
| MM105 | 1 | 1 | 3 | 4 | 1 | 1 | 1 | + | + | - | - | - | - | - | 1 | 1 | 1 | 1 | 1 | - | 1 | - |
| MM106 | 2 | 2 | - | 2 | 2 | 2 | 2 | + | + | 2 | 2 | - | 2 | 2 | 1 | 1 | 1 | 1 | 1 | - | 1 | 1 |
| MM107 | - | - | - | 1 | - | - | - | - | - | - | - | - | - | - | - | - | - | - | 3 | 2 | 1 | 1 |
| MM108 | 1 | 1 | 3 | 4 | 1 | 1 | 1 | + | + | - | - | - | - | - | 2 | 1 | 1 | 1 | 3 | 1 | 1 | 1 |
| MM109 | 1 | 1 | 2 | 3 | 2 | 2 | 2 | + | + | 2 | 1 | - | 1 | 2 | 1 | 1 | 1 | 1 | 2 | - | 1 | - |
| MM111 | 2 | 2 | 2 | 4 | 2 | 2 | 2 | - | + | 1 | 1 | - | 1 | 1 | 1 | 1 | 1 | 1 | 1 | 1 | 1 | 1 |
| MM117 | 1 | 1 | 1 | 2 | 1 | 1 | 1 | + | + | - | - | - | - | - | 2 | 1 | 1 | 1 | 1 | 1 | 1 | - |
| MM122 | 2 | 2 | 2 | 4 | 3 | 3 | 3 | + | + | - | - | - | - | - | 1 | 1 | 1 | 1 | 1 | 3 | - | - |
| MM128 | 1 | 1 | 1 | 2 | 1 | 1 | 1 | - | + | - | - | - | - | - | 1 | 1 | 1 | 1 | 2 | 2 | 1 | 1 |

Abbreviations: CA\_sulf, sulfate transporter family protein in cluster with carbonic anhydrase; \* pseudogene

Urease activity has been recently linked to an activity of the zinc-containing enzyme – carbonic anhydrase (CA) (Achal and Pan 2011). While urease maintains an alkaline environment by generating ammonia, carbonic anhydrase would provide carbon dioxide for biomineralization through dehydration of carbonic acid (H_2_CO_3_) also produced during ureolysis (Fig. 1). It was recently demonstrated that the activities of CA and urease were correlated along the bacterial growth and corresponded to maximum calcite production (Banks et al. 2010; Achal and Pan 2011), while inhibition of carbonic anhydrase activity decreased the rate of calcification, with calcite precipitation occurring more efficiently with the synergistic action of both enzymes – CA and urease (Dhami et al. 2014).

The presence of this highly efficient enzyme in cave settings was previously confirmed through metagenomic study of a speleothem in Tjuv-Ante’s Cave (Sweden) (Mendoza et al. 2016), and highly acidic cave biofilms, known as ‘snottites’ (Jones et al. 2011). However, it has never been directly associated to a specific taxonomic group through genome-based approach. The only link was suggested by Cuezva et al. (2012), who proposed CA to be responsible for CO_2_ sequestration by grey spot colonization found on the walls of Altamira cave, which were dominated by Actinobacteria.

Evaluation of the genomes of moonmilk phylotypes for the presence of β-CA (Smith and Ferry 2000), together with the sulfate transporter family protein in cluster with CA (SulP-type permease) revealed that all the investigated strains encode at least one copy of carbonic anhydrase (100% strains encoded β-CA (1) and 87% β-CA (2)) and 90% of them possessing sulfate transporter (Table 1). High copy number of this intracellular zinc metalloenzyme, reaching up to 7 copies in MM17, revealed their ubiquitous distribution among karstic bacteria, and suggests their applications in multiple and essential cellular processes beyond their presumed role in carbonate precipitation. Interestingly, a high representation of sulfate permeases of SulP family was also characteristic for the studied population. These broad specificity inorganic anion transporters were suggested to assist bicarbonate (HCO_3_^−^) transport (Felce and Saier 2005), which was experimentally confirmed in marine cyanobacteria (Price et al. 2004). While in cyanobacteria SulP transporter mediates HCO_3_^−^/Na^2+^ symport, the substrate specificity in the case of *Streptomyces* is highly speculative. Nevertheless, fusion of this protein with carbonic anhydrases indeed strongly suggests its participation in HCO_3_^−^ transport. However, whether it is an importer or exporter remains an open question. It could be possible that HCO_3_^−^ uptake through SulP increases the intracellular pool of this inorganic carbon species, which potentially might be efficiently transformed into carbon dioxide by carbonic anhydrases, and be exported outside the cell, unless required for cellular metabolism. Alternatively, SulP-dependent export of intracellularly formed HCO_3_^−^ through CA-mediated hydration of carbon dioxide, would be assisted with import of other ions (Fig. 1). Altogether, these findings reveal abundant distribution of genes involved in the inorganic ionic transport and metabolism, which might be related to the biomineralization phenomenon.

### Ammonification

Moonmilk *Streptomyces* were also evaluated for their ability to raise the pH through peptide/amino acid mineralization. 94 % of tested strains efficiently decomposed nitrogenous compounds into ammonia (Fig. 1, Fig. 3b, Fig. 4 and supplementary Fig. 4b). The majority displayed a strong metabolic phenotype (Fig. 5), which is unsurprising given that *Streptomyces* are well known ammonifying bacteria in soils, where they actively participate in the decomposition of organic matter (Prakash et al. 2012). In the isolated cave environment, with limited organic matter input from the surface, the source of such macromolecules is unclear, although peptides and amino acids might be entering the cave through water that has percolated through the soil, making such molecules more readily available than urea (Northup and Lavoie 2001). Amino acid/peptide ammonification was found to be more widespread among the moonmilk isolates than ureolysis, supporting this hypothesis.

**Fig. 5.**
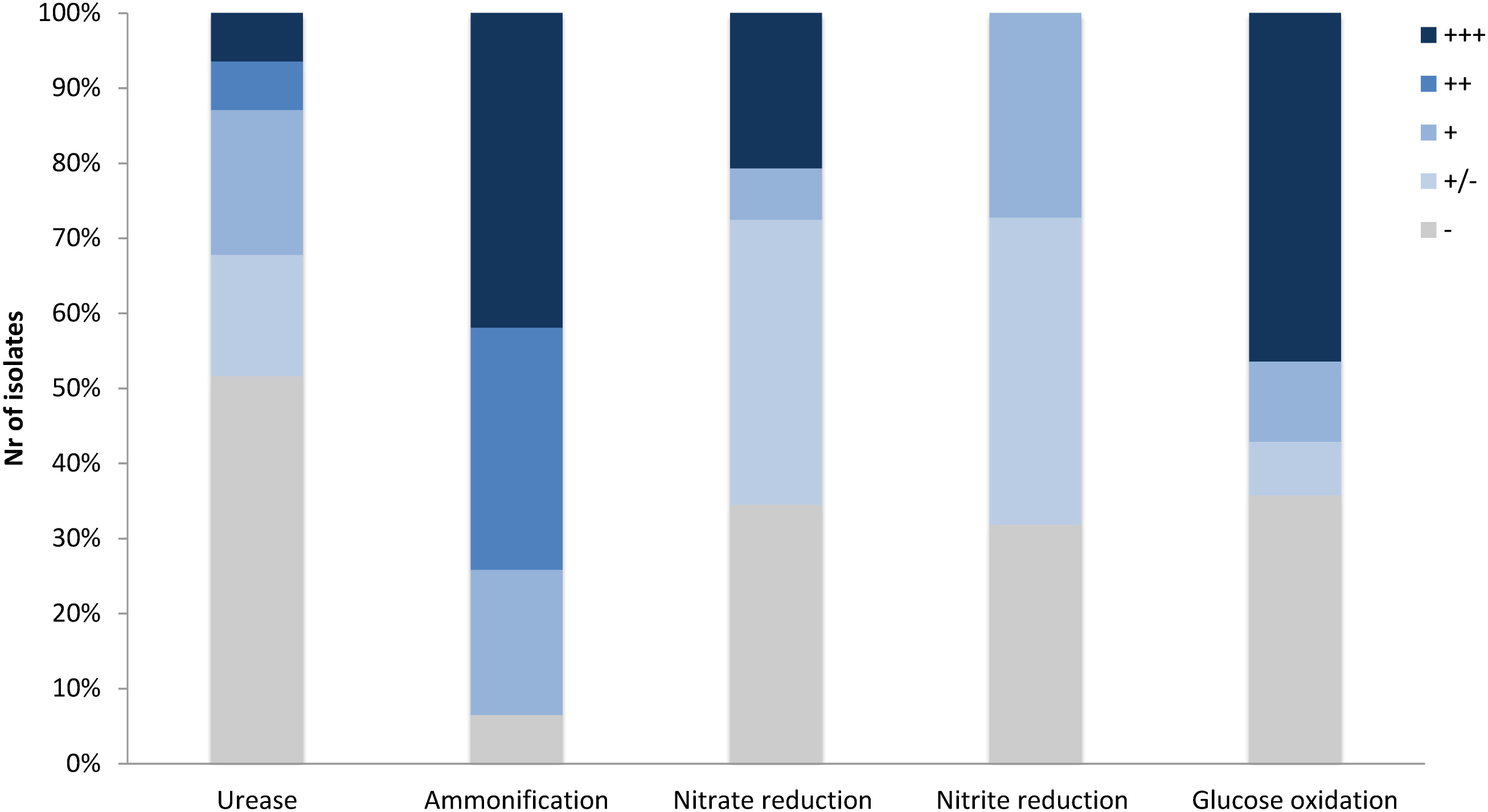
Percentage of moonmilk *Streptomyces* displaying carbonatogenesis related activities *in vitro* within different metabolic categories. (-) lack, (+/-) weak, (+) moderate, (++) good, and (+++) strong

### Dissimilatory nitrate reduction to ammonium

Dissimilatory nitrate reduction to ammonium (DNRA) is another nitrogen cycle-related process considered to be involved in calcification (Castanier et al. 2000) (Fig. 1). This pathway operates in oxygen-limited environments, which could be encountered in the inner layers of the moonmilk deposits. Moreover, moonmilk develops alongside dripping water and its often pasty structure can become fluid, based on its water content, which can drastically reduce oxygen availability. Although *Streptomyces* are obligate aerobes, they are genetically capable of survival under oxygen-limited conditions, encoding genes related to anaerobic respiration, the so-called “anaerobic paradox” (Borodina et al. 2005). Indeed, the model species *Streptomyces coelicolor* was reported to anaerobically respire nitrate (Fischer et al. 2010; Fischer et al. 2014). Though nitrite, the product of this process, was not reduced to ammonia, but detoxified through extrusion via NO_3_^−^/NO_2_^−^ antiporter system (Fischer et al. 2010; Fischer et al. 2012). In *S. coelicolor* the NirBD reductase was demonstrated to participate in nitrogen assimilation; however a *nirBD* null-mutant grown in the presence of nitrite and excess ammonium was still able to reduce nitrite suggesting the activity of an alternative and yet unknown enzyme (Fischer et al. 2012). We therefore questioned whether moonmilk isolates would be able to mediate DNRA and reduce nitrate and nitrite under oxygen-limited conditions. For this purpose, we incubated each strain in static (without agitation) liquid culture conditions to limit oxygen availability. Overall, 68% and 77% of strains revealed a capacity to reduce either nitrate (NO_3_^−^) or nitrite (NO_2_^−^), respectively, with 52% able to reduce both nitrate and nitrite (Fig. 3c/d, Fig. 4 and Fig. 5). No N_2_ gas production was observed, excluding denitrification, which has been only rarely reported for *Streptomyces* (Albrecht et al. 1997; Shoun et al. 1998; Kumon et al. 2002). The lack of N_2_ generation would suggest a complete reduction of nitrate/nitrite to ammonia via dissimilatory nitrate reduction. Ammonia produced by this pathway, unless not incorporated by other bacteria or oxidized to other nitrogenous compounds, might alkalinize the extracellular environment and thus stimulate CaCO_3_ precipitation (Fig. 1). In order to evaluate the presence of a DNRA pathway in moonmilk isolates we screened their genomes for the presence of genes coding for respiratory nitrate reductases (*narGHJI*) and their associated NarK-type nitrate/nitrite transporter - NarK2 (Fig. 1). While 40% of moonmilk strains encoded NarK2 transporter, 30% of them possessed the respiratory nitrate reductases genes (*narGHJI*), with isolates MM7, MM10, MM48, MM106, MM109, MM111, encoding a complete *nar* cluster (Table 1). Most of those strains were found to be among the strongest nitrate reducers under oxygen-limited conditions (Fig. 4). The fact that only a minority of the moonmilk *Streptomyces* possessed the genetic material to perform the first step of DNRA, while a majority (68%) was able to reduce nitrate, suggests that another nitrate reduction pathway was operating under the condition tested. This alternative pathway is most likely the NO_3_^−^ and NO_2_^−^ assimilatory process that uses nitrate and nitrite as nutrient sources through assimilatory *nasA* (nitrate reductase) and *nirBD* (nitrite reductase) genes, with the assistance of additional Nark-type NO_3_^−^ transporter (Tiffert et al. 2008; Amin et al. 2012; Fischer et al. 2012). Assimilatory reduction of nitrate and nitrite to ammonium is highly plausible as 93% of moonmilk strains possess *nasA*, *nirBD*, and *narK* NO_3_^−^ importer genes (Table 1). When monitored on solid medium and thus without oxygen limitation, 68% and 77% of the tested strains were able to reduce nitrate and nitrite, respectively (supplementary Fig. 4c/d), which confirmed the high potential of moonmilk *Streptomyces* to use nitrate and nitrite as a nitrogen source, as previously reported for terrestrial *Streptomyces* (Pullan et al. 2011; Fischer et al. 2012). Altogether, although moonmilk *Streptomyces* possess metabolic ability to reduce nitrate and nitrite, without production of gas, pointing on ammonia as a final product, we cannot at this point conclude whether this process represents assimilatory or dissimilatory pathway and whether DNRA is fully functional.

### Active calcium transport

In addition to processes passively influencing carbonate precipitation, bacteria can also actively impact this phenomenon, through an active transport of calcium ions. Banks et al. (2010) suggested that the calcium-toxicity driven removal of this ion outside the bacterial cell is a factor driving calcification phenotype. Therefore, we have also retrieved through an *in silico* search ChaA, the Ca^2+^/2H^+^ antiporter system suggested to be involved in CaCO_3_ deposition (Hammes and Verstraete 2002; Banks et al. 2010) (Fig. 1). The presence of the *chaA* gene was confirmed for 50% of moonmilk cultivable phylotypes (Table 1), extending in those strains the calcium-detoxification system to their spectrum of biomineralization-related processes.

### Production of CaCO_3_ deposits by moonmilk *Streptomyces*

In order to confirm whether the moonmilk cultivable *Streptomyces* could indeed produce mineral deposits, we selected two phylotype representatives to be first investigated by polarized light microscopy then by ESEM in low vacuum mode for the presence of calcium carbonate precipitates. The selection of strains was based on their predispositions for CaCO_3_ precipitation as judged by the sum of metabolic performance observed for ureolysis and peptide/amino acid ammonification – the two most significant activities observed for moonmilk *Streptomyces* (Fig. 4). Strains MM24 and MM99, amongst one of the best isolates based on the metabolic ranking (Fig. 4), were simultaneously cultivated on urea-containing CPA medium for two months, as well as for one month on the modified B-4 medium commonly used for CaCO_3_ precipitation assays. Combined microscopic observations of bacterial colonies surfaces with BSE and GSE detectors revealed highly abundant calcite deposits produced by both strains in both media tested (Fig. 6 and Fig. 7). Ureolysis-mediated mineral precipitation was confirmed by the lack of any calcite in urea-deficient CPA medium for both isolates (data not shown). All the produced mineral deposits appeared bright under polarized light (data not shown). The morphology of the calcite polymorphs was comparable between the isolates, however differed between the two culture conditions. On BSE-images, bacterial colonies grown in CPA medium showed a rocky surface with discoidal- or oval-shaped structures of variable diameter that were almost completely encrusting microbial colonies (Fig. 6a/c). The mineral nature of the deposits was confirmed through their high (white) contrast on BSE-images compared to the surrounding dark organic matter of the colonies (Fig. 6a/c). In addition, their CaCO_3_ mineral composition was confirmed by the elemental X-ray analyses (supplementary Fig. 5) and elemental mapping (Fig. 6b/d). Calcium, carbon and oxygen were present roughly in stoichiometric proportion of CaCO_3_ in the spectra (supplementary Fig. 5), and the distribution of those elements was clearly visualized on the mapping (Fig. 6b/d). The presence of calcium was associated with CaCO_3_ minerals, while higher proportion of carbon was associated with organic colony biomass (Fig. 6b/d). Interestingly, just on the mineral surface, dense webs of nano-sized filaments were observed (Fig. 7a-e). On high resolution BSE-images, they appeared either as dark curved filaments at the mineral surface or in the middle of mineralized wrinkles suggesting that bacteria produced minerals in which they got progressively entombed (Fig. 7a-d). The filaments free of mineral were also clearly seen interconnecting together and connecting adjacent mineral deposits (Fig. 7e). While on the CPA medium both strains prolifically produced relatively small-sized calcite polymorphs (up to 100 µm) (Fig. 6a/c), on the B-4 medium the observed precipitates although being more scarce, were much larger (up to 400 µm) (Fig. 6e/g). The inorganic nature of the larger CaCO_3_ deposits (Fig. 6e/g) were confirmed by X-ray elemental spectra (supplementary Fig. 5) and elemental mapping (Fig. 6f/h), which clearly distinguished CaCO_3_ minerals from the surrounding bacterial biomass rich in carbon and oxygen. On BSE-image (Fig. 7f) and under polarized light (data not shown), tiny CaCO_3_ deposits were also detected along randomly distributed bacterial filaments. Additionally, unlike in CPA medium where the mineral surface was rather irregular and unstructured, mineral produced by both strains in B-4 showed morphologically distinct areas, either with a smooth, radiating texture (Fig. 7h), with visible filamentous imprints (Fig. 7j) or wrinkles (Fig. 7g), that presumably corresponded to bacterial nano-sized filaments.

**Fig. 6.**
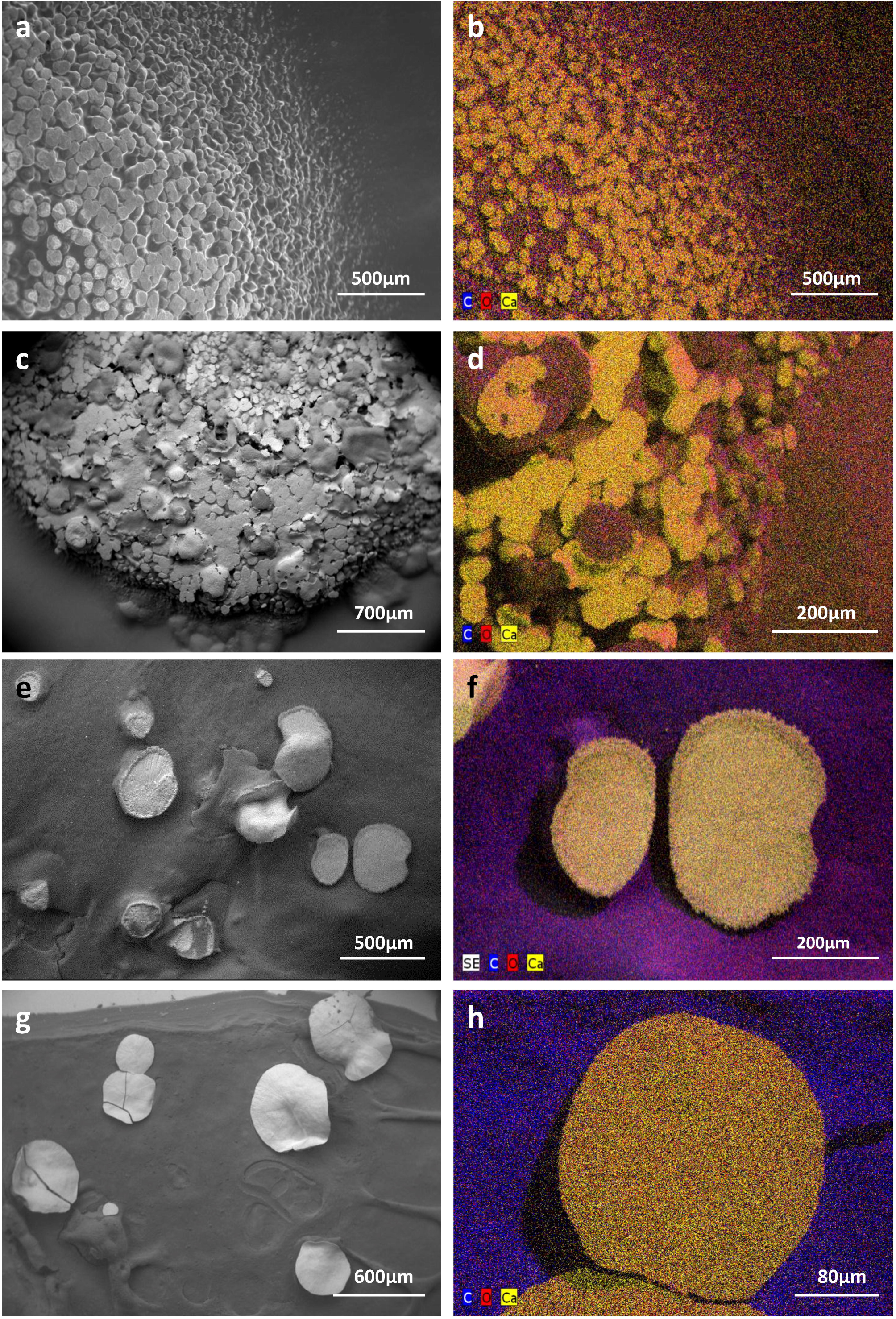
Low vacuum-SEM-BSE images (a, c, e, g) and X-ray elemental mappings (b, d, f, h) of CaCO_3_ deposits produced by MM24 (a-b, e-f) and MM99 (c-d, g-h) isolates. Abundant mineral deposits produced by isolates MM99 **(a)** and MM24 **(c)** in CPA medium, nearly encrusting the whole microbial colony, were morphologically different from less abundant, but much bigger, mineral polymorphs produced in modified B-4 agar by MM99 **(e)** and MM24 **(g)**. The observed precipitates were found to be CaCO_3_ minerals as revealed by elemental spectra (supplementary Fig. 5) and mappings in both media and for both isolates (MM99 **(b)** and MM24 **(d)** in CPA, MM99 **(f)** and MM24 **(h)** in modified B-4). The mappings for MM99 and MM24 in B-4 are marked with the white squares. The mappings are combined images in which detected dominant elements, namely carbon (C), oxygen (O), and calcium (Ca) are assigned to a defined color

**Fig. 7.**
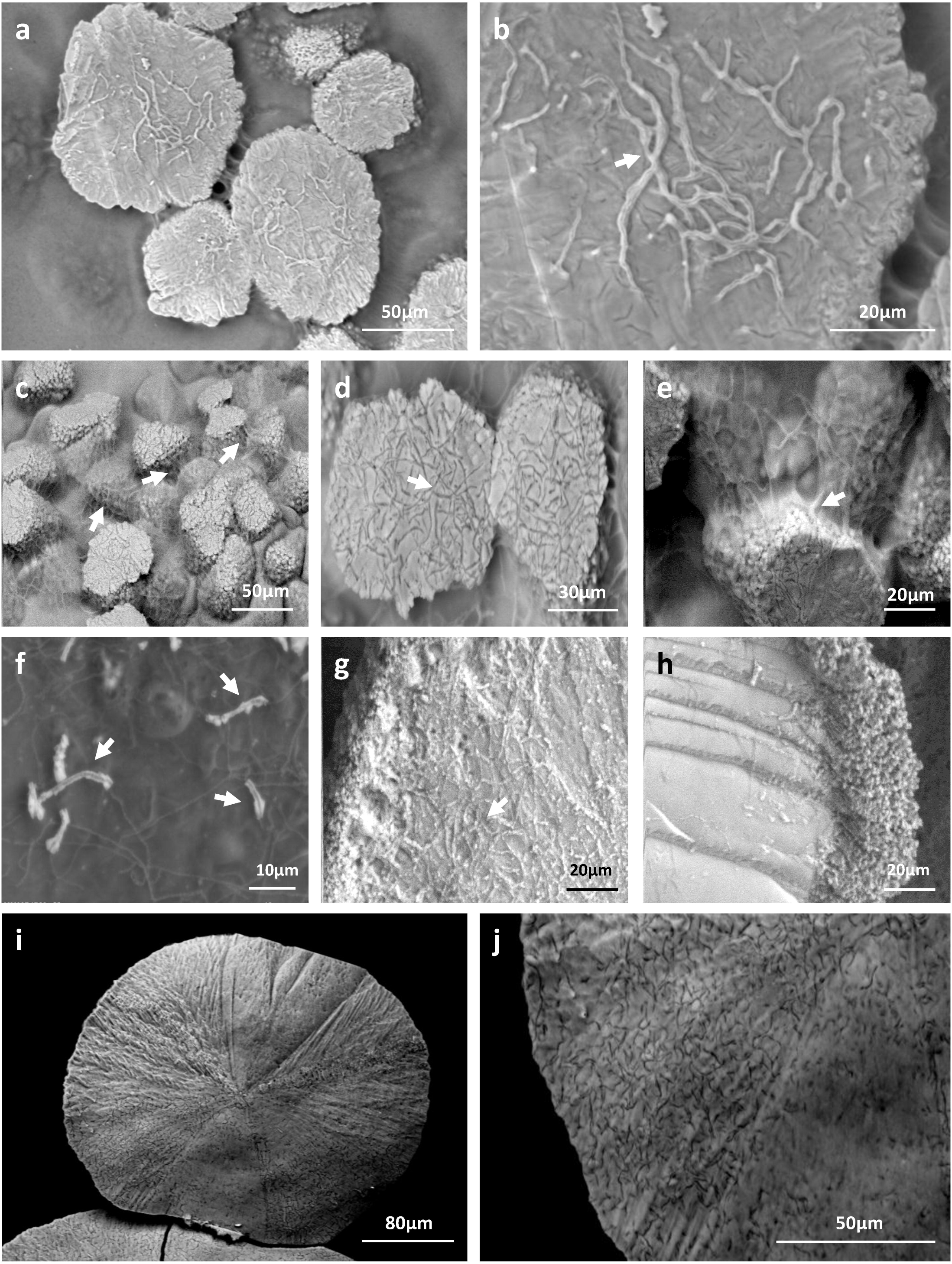
Biogenic signatures within mineral deposits produced by moonmilk-originating *Streptomyces* (LV-SEM-BSE images). Along with the different morphologies of produced mineral deposits specific to the culture conditions used, numerous bacterial imprints were observed. *Streptomyces* filaments were completely incorporated into the produced minerals as observed for MM99 **(a/b)**, MM24 **(c/d/e)** in CPA medium, as well as for MM99 **(g)** and MM24 **(i/j)** in modified B-4 agar. Their mineralization also appeared as wrinkles each containing a bacterial filament suggesting their progressive encrustation within the mineral **(b/e/g**, arrows). Alternatively, non-mineralized microbial filaments were seen interconnecting adjacent mineral deposits, as detected for MM24 in CPA **(c/e**, arrows), or possible compacted aggregates of filaments were observed at the edges of the mineral produced by MM99 in B-4 **(h)**. Initial steps of mineral deposition were detected along bacterial filaments in modified B-4 culture of isolate MM99 **(f**, arrows)

### Potential role of cultivable moonmilk-derived *Streptomyces* in carbonate dissolution

In addition to constructive processes, bacteria are also believed to induce cave bedrock dissolution. As oppose to precipitation, a dissolution phenomenon is related to the acidification of the bacterial microenvironment, most likely as a result of organic acid production, which are the by-products of microbial carbon metabolism. The presence of detectable levels of organic acids in cave environment was previously demonstrated by *in situ* analysis via ATR-FTIR spectroscopy (Bullen et al. 2008). Released organic acids are able to bind cations such as Ca^2+^ and liberate carbonates, which can be subsequently re-precipitated to form cave secondary deposits or be used by bacteria. Therefore, we have tested the 31 phylotypes for their ability to decrease the pH of the medium through oxidative degradation of glucose based on the standardized oxidative/fermentative test (Hugh and Leifson 1953). Over 68% of isolates were found to induce media acidification by this pathway, which was observed as yellow color to transparent halo *503* development around the inoculum (Fig. 8a and Fig. 9). Among them, 52% of the *Streptomyces* strains exhibited either good or strong oxidative glucose respiration abilities, strongly reducing the extracellular pH (Fig. 5, Fig. 9 and supplementary Fig. 6).

**Fig. 8.**
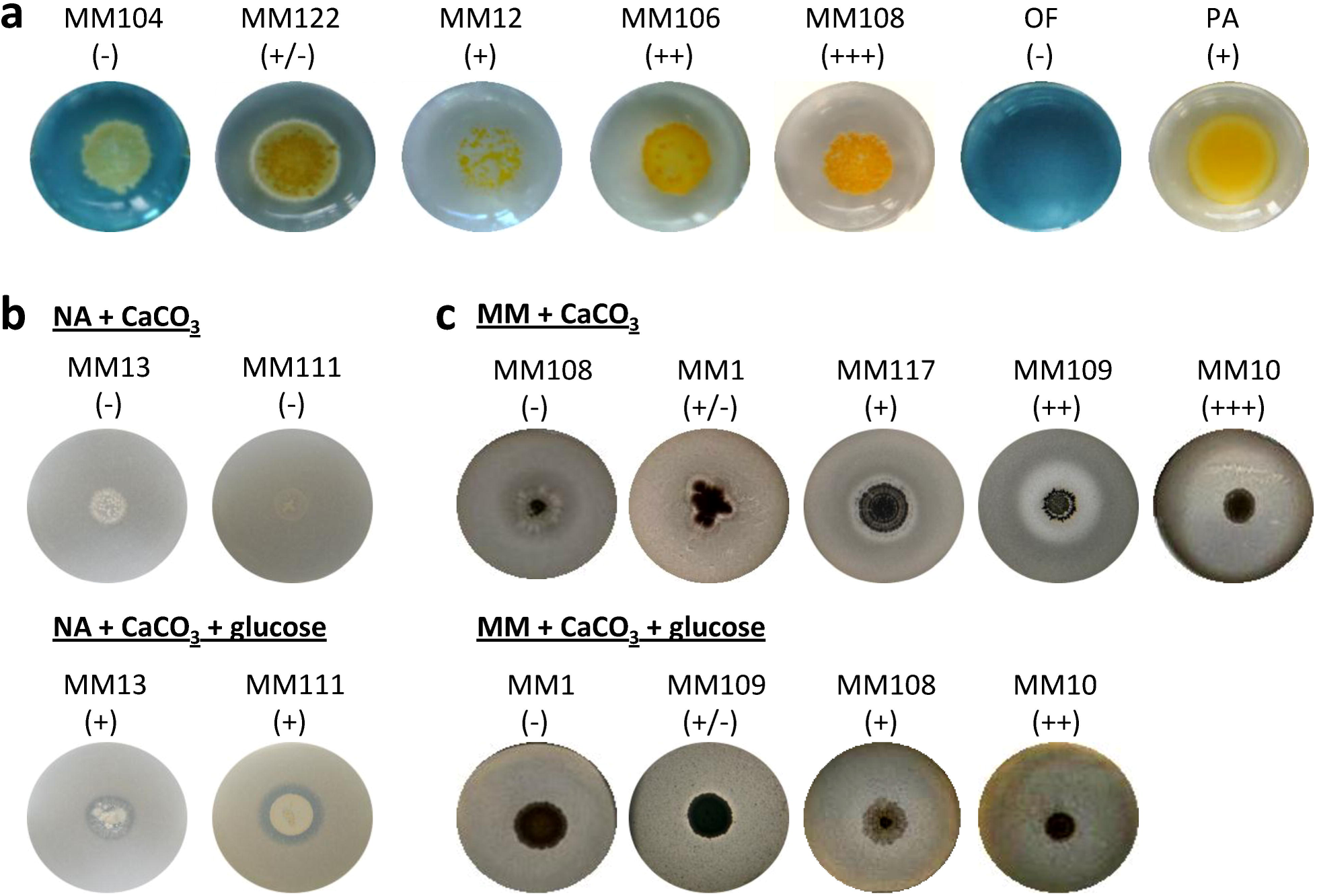
Dissolution-related metabolic activities of the moonmilk *Streptomyces*. Metabolic performance observed for oxidative glucose degradation **(a)**, and calcium carbonate dissolution in diluted nutrient agar **(b)** or minimal media **(c)**. *Pseudomonas aeruginosa* ATCC 27853 (PA) was used as positive control in glucose oxidative assay. The metabolic performance is designated through the symbols: (-) lack, (+/-) weak, (+) moderate, (++) good, and (+++) strong activity. Media not supplemented with glucose are in the upper line in panel **b** and **c**, while media with addition of glucose are displayed in the bottom line of the corresponding panels

**Fig. 9.**
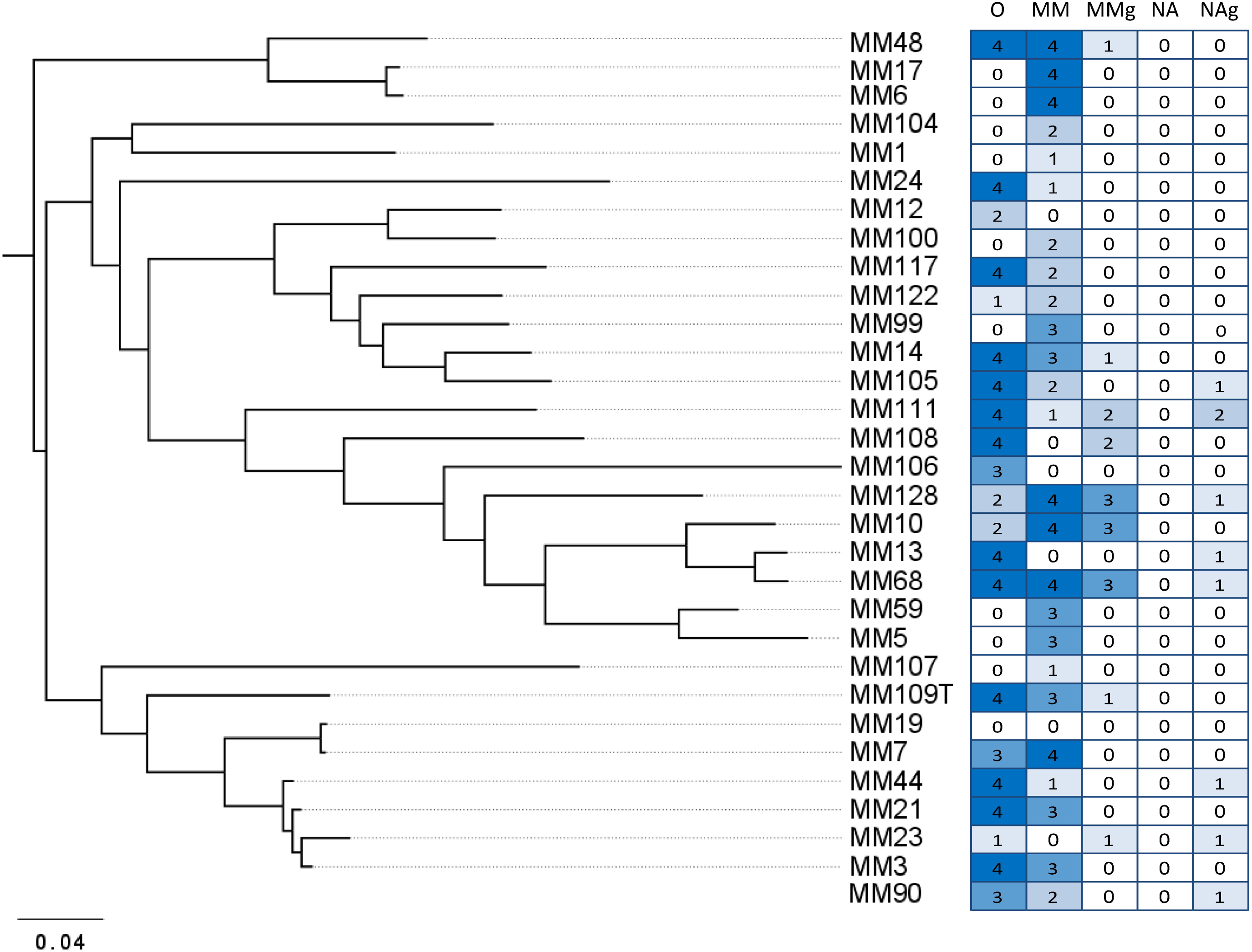
The global pattern of carbonate dissolution-associated metabolic activities of moonmilk *Streptomyces.* The heatmap plot representing metabolic performance of moonmilk *Streptomyces* strains indicated by color scale, correlated with their MLSA-based phylogenetic classification (Maciejewska et al. 2016). Abbreviations: O, glucose oxidation; MM, minimal media with CaCO_3_; MMg, minimal media with CaCO_3_ and glucose; NA, diluted nutrient agar with CaCO_3_; NAg, diluted nutrient agar with CaCO_3_ and glucose

We have further tested solubilization abilities related to carbon metabolism by cultivating the phylotypes in calcium carbonate containing media – either in minimal medium or in diluted nutrient agar - supplemented or not with glucose. The clear effect of glucose breakdown was observed in diluted nutrient agar in which the supplementation with the carbohydrate induced CaCO_3_ dissolution in 26% of the isolates, while none of the phylotypes promoted dissolution in the non-supplemented medium (Fig. 8b and Fig. 9). In the non-supplemented nutrient agar, rich in organic nitrogen source, the cellular energy comes from amino acid utilization, releasing ammonia as the by-product (Fig. 1), which increases the pH of the medium and thus promotes precipitation rather than dissolution. The addition of glucose clearly induces the opposite effect but only in a minority of the isolates which suggests the preference towards amino acids as carbon source over glucose in the large majority of the studied strains (Fig. 8b and Fig. 9). On the contrary, when assays were performed in the minimal medium, CaCO_3_ dissolution was instead rather inhibited by the exogenous supply of glucose, as only 29% of isolates showed ability to solubilize CaCO_3_ under this condition, while in minimal media without glucose supply the dissolution phenotype was characteristic for 81% of isolates (Fig. 8c and Fig. 9). This might be probably related to the fact that in minimal medium not supplemented with glucose the only carbon source constitutes the carbonate/bicarbonate from CaCO_3_, which is probably efficiently scavenged by cave-dwelling bacteria for autotrophic growth resulting in the high dissolution rate observed under this condition. Although *Streptomyces* are mainly heterotrophic microorganisms, autotrophic growth using CO or CO_2_ as a sole carbon source within members of this genus has already been reported (Kim et al. 1998; Gadkari et al. 1990). The identification of a high number of carbonic anhydrases together with SulP transporters among moonmilk *Streptomyces* (Table 1) could propose a mechanism via which extracellular bicarbonate would be incorporated into the cell and subsequently converted to CO_2_ (Figure 1). In the moonmilk niche that is deprived of organic carbon, the uptake of inorganic carbon could thus be a possible scenario, which would primarily promote CaCO_3_ precipitation, and in a second step would lead to CaCO_3_ dissolution, as a result of organic acids excretion. The availability of glucose also induces a dissolution phenotype via release of organic acids from glucose breakdown, however only in 3 out of the 31 strains tested. This indicates that, although carbohydrate metabolism might somehow play a role in rock weathering, it is probably not the only operating system leading to this phenomenon, particularly in carbon-limited cave environment, where the source of carbon might be the rock itself.

## Discussion

If Actinobacteria really participate in the genesis of moonmilk deposits which metabolic activities would potentially be involved? This was the main question addressed in our study which used a collection of *Streptomyces* strains isolated from moonmilk in order to provide metabolic and genetic evidences of their presumed role in mediating the formation of these speleothems. Metabolic profiling revealed that all of the isolated *Streptomyces* possessed the capacity to promote calcification through at least one pathway involved in biomineralization. Ammonification of peptides/amino acids was found to be the most widespread and the strongest activity. This could be in agreement with the fact that peptides and amino acids can constitute self-sustainable sources of carbon and nitrogen for bacteria and thus support growth of microbial populations irrespectively of allochtonous nutrient input. Interestingly, genome mining extended the possible spectrum of metabolic capacities as it revealed the presence of additional pathways in each phylotype involved in biomineralization processes, either related to CO_2_ hydration or active transport of calcium ions. These findings, supported by microscopy observations of bacteria-like filaments in moonmilk deposits and crystals produced by individual moonmilk-originating bacteria, confirm its biogenic origin and the importance of filamentous Actinobacteria in its genesis. However, the metabolic activities evaluated *in vitro* were not always directly related to the genetic predisposition of individual strains as some isolates with great genetic potential remained metabolically silent. This may suggest that they were grown under conditions too different from those encountered in their original niche to trigger the investigated activity or that specific environmental cues are required for their activation. Consequently, whether those processes are indeed active *in situ* also remains an open question. Additionally, our collection of moonmilk Actinobacteria, though being the most significant population from these speleothems generated so far, does not include the large majority of endemic representatives which are viable but not cultivable A metaproteomic analysis of proteins extracted from moonmilk deposits is most likely the only approach that would accurately identify the strains that importantly participate in carbonatogenesis and the metabolic pathways involved. This approach is currently under investigation.

## Acknowledgments

MM and LM work is supported by a Research Foundation for Industry and Agriculture (FRIA) grant. AN work is supported by a First Spin-off grant from the Walloon Region (Grant number: 1510530; FSO AntiPred). Computational resources (“durandal” grid computer) were funded by three grants from the University of Liège, “Fonds spéciaux pour la recherche,” “Crédit de démarrage 2012” (SFRD-12/03 and SFRD-12/04) and “Crédit classique 2014” (C-14/73) and by a grant from the F.R.S.-FNRS “Crédit de recherche 2014” (CDR J.0080.15). This work is supported in part by the Belgian program of Interuniversity Attraction Poles initiated by the Federal Office for Scientific Technical and Cultural Affairs (PAI no. P7/44). The authors thanks Isabelle Habsch for microscopy sample preparation and the Centre of Aid for Research and Education in Microscopy (CAREm-ULg) for giving access to SEM-equipment. SR and MH are Research Associates at Belgian Fund for Scientific Research (F.R.S-FNRS).

